# Developmental and molecular contributions to contextual fear memory emergence in mice

**DOI:** 10.1101/2023.02.03.527024

**Authors:** Alexandra L. Lanjewar, Pat Levitt, Kathie L. Eagleson

## Abstract

Cognitive impairment is a common phenotype of neurodevelopmental disorders, but how these deficits arise remains elusive. Determining the onset of discrete cognitive capabilities facilitates studies in probing mechanisms underlying their emergence. The present study analyzed the emergence of contextual fear memory persistence (7-day memory retention) and remote memory (30-day memory retention). There was a rapid transition from postnatal day (P) 20 to P21, in which memory persistence emerged in C57Bl/6J male and female mice. Remote memory was present at P23, but expression was not robust compared to pubertal and adult mice. Previous studies reported that following deletion of the MET receptor tyrosine kinase (MET), there are fear memory deficits in adult mice and the timing of critical period plasticity is altered in the developing visual cortex, positioning MET as a regulator for onset of contextual fear memory. Sustaining *Met* past the normal window of peak cortical expression or deleting *Met*, however, did not alter the timing of emergence of persistence or remote memory capabilities during development. Fear memory in young adults, however, was disrupted. Remarkably, compared to homecage controls, the number of FOS-expressing infragranular neurons in medial prefrontal cortex (mPFC) did not increase from contextual memory formation recall of fear conditioning at P35 but exhibited enhanced activation at P90 in male and female mice. Additionally, MET-expressing neurons were preferentially recruited at P90 compared to P35 during fear memory expression. The studies demonstrate a developmental profile of contextual fear memory capabilities. Further, developmental disruption of *Met* leads to a delayed functional deficit that arises in young adulthood, correlated with an increase of mPFC neuron activation during fear memory recall.

## INTRODUCTION

Cognitive development is a protracted process, and brain regions involved, such as medial prefrontal cortex (mPFC), continue to mature into early adulthood [1-5], coinciding with increased and optimized cognitive capacity [6-8]. Cognitive deficits are common in neurodevelopmental disorders (NDDs) [9-15]; however, how and when such deficits arise remain elusive.

In rodents, contextual fear paradigms are used to assess learning and memory functions, which involve context encoding, fear encoding, and associative learning and memory. More specifically, for contextual fear memory to occur, the hippocampus encodes contextual information of the conditioning, the amygdala encodes the emotional properties of the context, while the mPFC encodes both contextual and fear processing [16]. Perturbations in the expression of NDD risk genes during development can lead to contextual fear deficits in mice. Most studies, however, focus analyses on adults [17-19], leaving a knowledge gap in the temporal and mechanistic origins of cognitive deficits. Studies have begun to focus on the normative ontology of contextual fear learning abilities in wild type mice. Akers et al. reported that as early as postnatal day (P) 15, mice form a fear memory that lasts at least 1-day (d) but does not persist for 7-d [20]. By P25, longer-term memory capabilities are present, such that mice can retain fear memories for at least 30-d [21]. These studies identify a broad window of cognitive development, during which longer-term memory capabilities arise. The precise temporal trajectory, however, remains unknown.

Accumulating evidence underscores a key role for the MET receptor tyrosine kinase (MET) in the development of discrete circuits within the forebrain. In cerebral cortex, abundant MET expression coincides with the period of peak synaptogenesis, and MET signaling modulates dendritic and synapse development and maturation [22-27]. Prolonging or eliminating MET expression alters the timing of critical period plasticity for ocular dominance, shifting the critical period later or earlier, respectively [25]. Additionally, prolonging MET expression during a critical period for social cognition alters social behavior in adult mice [26]. Finally, adult mice in which *Met* had been conditionally deleted embryonically in all neural cells or in cells arising from the dorsal pallium exhibit contextual fear learning deficits [28-30]. Thus far, no study has determined when these deficits arise. Together, these studies position MET as a candidate for regulating the timing of onset for contextual fear memory capabilities.

Here, we determine the precise developmental trajectory for onset of 7-d persistent memory capabilities and whether this coincides with the onset of remote memory capabilities (30-d memory) [31-32]. We also determine whether sustaining or eliminating MET signaling affects contextual fear memory abilities prior to adulthood. Lastly, experiments were performed to determine the developmental and adult activity of MET^+^ neurons in mPFC during contextual fear memory.

## MATERIALS AND METHODS

### Animals

Mice were bred in the Children’s Hospital Los Angeles (CHLA) vivarium and housed on ventilated racks with a 13:11 hour light:dark cycle at 22°C with ad libitum access to water and a standard chow diet (PicoLab Rodent Diet 20, #5053, St. Louis, MO). All mouse lines were maintained on a C57Bl/6J background. Therefore, the C57Bl/6J strain (The Jackson Laboratory) was used as the wild type (WT) mouse line to determine the normal developmental trajectory for retaining a 7-d persistent memory or a 30-d remote memory. To sustain MET expression in all dorsal pallial excitatory neurons, a <u>c</u>ontrollable transgenic <u>o</u>verexpression for *<u>Met</u>*(cto-*Met*) mouse line was used, as previously described and validated [26]. Sustained MET expression in the cerebral cortex of the cto-*Met* mouse line was validated in our laboratory by Western blot to ensure consistent inter-lab colony transgene expression (data not shown). In this line, the *Met* transgene is expressed abundantly by P16 under the control of the CAMKII promoter, resulting in increased MET signaling [25-26]. Thus, MET expression is sustained throughout the course of the experiments. Littermates not expressing the *Met* transgene were considered controls. A mating scheme that has been previously validated in the lab was used to reduce or delete the *Met* gene by pairing *Met^fx/fx^* females and *Nestin^cre^*; *Met^fx/+^* males to produce control (*Met^fx/fx^*: Cre-negative;*Met^fx/fx^* or *Met^fx/+^*: Cre-negative;*Met^fx/+^*), conditional heterozygous (cHet: *Nestin^cre^;Met^fx/+^*), and conditional homozygous (cKO: *Nestin^cre^;Met^fx/fx^*) pups [28]. This ensured the production of sufficient numbers of the control and mutant genotype combinations in each litter to reduce the potential confound for inter-litter variability in the fear conditioning paradigm. Since there were no differences in freezing behavior in *Met^fx/fx^* and *Met^fx/+^* mice (data not shown), data were collapsed into a single control group that produces normal MET levels. Importantly, a previous study reported no genotype effect of aberrant locomotor behavior in the Nestin-Cre line when *Met* is conditionally deleted or reduced [28], so our reported freezing responses are independent of a locomotion difference in these mice. The cHet mice produce 50% of normal MET levels and the cKO mice do not produce any MET in neural cells [28-29]. A separate breeding scheme - WT females crossed with *Nestin^cre^*males - was used to produce Cre+ or Cre- mice to determine whether the *Nestin^cre^*driver alone results in developmental fear memory deficits, as previously reported for adult males [33]. Lastly, a *Met*^EGFP^ BAC transgenic mouse line was used to visualize green fluorescent protein (GFP) in MET-expressing neurons in mice homozygous for the *Met-EGFP* transgene (*Met*^GFP^) [34-35]. For all mice, the day of birth was designated P0. At P21 (+/-1 d), mice were weaned and housed with same-sex littermates (2-5/cage). To address potential litter and sex effects, each experimental group included a maximum of 2 males and 2 females from a single litter, with a minimum of three litters and approximately equal numbers of males and females represented. For all analyses, investigators were blind to genotype and condition. Animal care and experiments conformed to the guidelines set forth by the CHLA Institutional Animal Care and Use Committee.

### Contextual Fear Conditioning and Testing

Contextual fear conditioning and memory testing were performed, following the Akers et al. paradigm that utilizes relatively low shock intensity and number [20] (Fig. 1A). The present study used the NIR Video Fear Conditioning Package for Mouse (MED-VFC2-USB-M; Med Associates Inc, Georgia, VT). Fear conditioning chambers (Med Associates VFC-008-LP) were 29.53×23.5×20.96cm with shock-grid floors (Med Associates VFC-005A). Shocks were generated by a standalone aversive stimulator/scrambler (Med Associates ENV-414S). Separate cohorts of mice from each mouse line were trained on P15, P20, P21, P22, P23, P35, P46-P53 (post-pubertal adolescents, denoted P50 from hereon) or P89-P99 (adults, denoted P90 from hereon). Briefly, “Shock” mice were acclimated in the chamber for 2-min and then presented with 3 unsignaled 2-sec foot shocks of 0.5mA intensity spaced 1-min apart. One minute after the last shock, mice were removed from the chamber and returned to their home cage. Shock delivery was confirmed by mice jumping and/or vocalizing during the shock. In the instances that shock delivery could not be confirmed for all three shocks, mice were excluded from further testing (<5%); there were no age, sex, or genotype differences in exclusions. Age-matched littermates designated “No-Shock” mice were placed in the chamber for the same length of time without receiving foot shocks and served as controls for spontaneous (non-memory induced) freezing. Memory trials were conducted 1- (formation), 7- (persistence), 14- (longer persistence), or 30- (remote) d later. On the testing day, mice were returned to the chamber and allowed to explore for 5-min (or 2-min in FOS experiments) without any foot shocks presented.

**Fig. 1:**
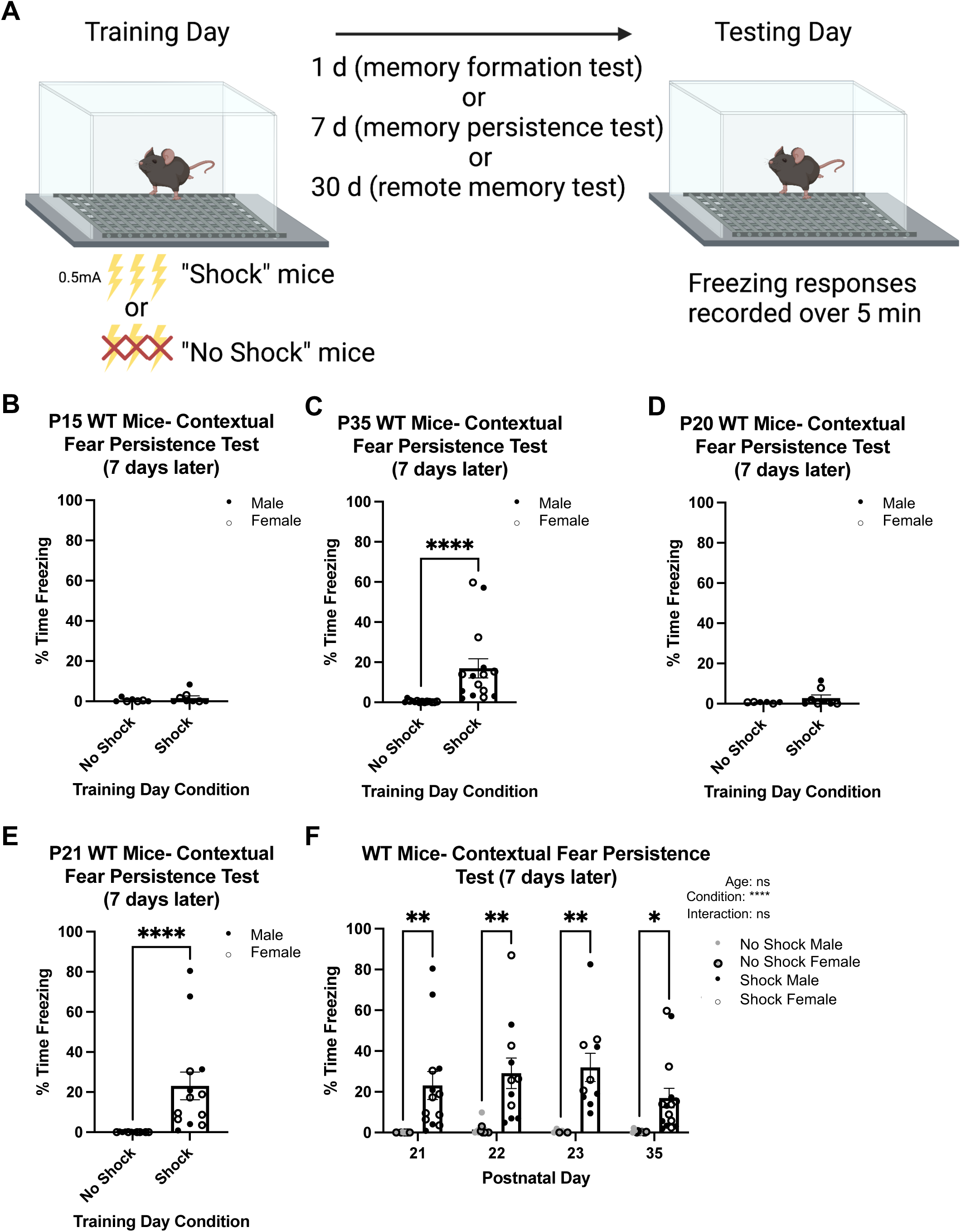
Contextual fear memory persistence rapidly emerges at P21 in WT mice. **A** Diagram of contextual fear memory persistence paradigm used in the present study. **B** Quantification of the percentage time freezing on testing day of No Shock (n = 7) and Shock (n = 8) mice trained on P15 and tested 7 d later. The Shock group did not exhibit significantly more freezing than the No Shock mice. **C** Quantification of the percentage time freezing on testing day of No Shock (n = 14) and Shock (n = 15) mice trained on P35 and tested 7 d later. The Shock group exhibited significantly more freezing than No Shock (\*\*\*\**p* < .0001). **D** Quantification of the percentage time freezing on testing day of No Shock (n = 6) and Shock (n = 8) mice trained on P20 and tested 7 d later. The Shock group did not exhibit significantly more freezing than the No Shock mice. **E** Quantification of the percentage time freezing on testing day of No Shock (n = 12) and Shock (n = 13) mice trained on P21 and tested 7 d later. The Shock group exhibited significantly more freezing than No Shock (\*\*\*\**p* < .0001). **F** Quantification of the percentage time freezing on testing day of No Shock and Shock mice trained at different ages and tested 7 d later (P21 No Shock: n = 12; P21 Shock: n = 13; P22 No Shock: n = 10; P22 Shock: n = 11; P23 No Shock: n = 6; P23 Shock: n = 10; P35 No Shock: n = 14; P35 Shock: n = 15). Shock mice exhibit significantly more freezing than No Shock mice at each (\**p* < .05; \*\**p* < .01, \*\*\*\**p* <.0001, ns, not significant).

### Behavioral Analysis

During testing day, freezing responses were recorded by a monochrome video camera (Med Associates VID-CAM-MONO-5). A freeze response was considered no movement above an 18au threshold for at least 1-sec (30 frames), analyzed by automated software (Video Freeze, SOF-843, Med Associates). The percentage of time freezing over the 5-min (or 2-min) trial on testing day was calculated by the automated software.

### Immunohistochemistry Staining

At P36 or P91, 90-min after testing of contextual fear memory formation, *Met*^GFP^ brains were collected, processed, and immunostained as described [36]. Primary antibodies (1:500) used were chicken anti-GFP (Abcam Cat#ab13970, RRID:AB_300798), rat anti-CTIP2 (Abcam Cat#ab18465, RRID:AB_2064130), and rabbit anti-CFOS (Cell Signaling Technology Cat#2250, RRID:AB_2247211). Alexa Fluor^®^ F(ab’)_2_ conjugated secondary antibodies (1:500; Abcam) were used.

### Imaging and Cell Count Analysis

Sections including mPFC (corresponding to areas 24a, 25, and 32 [37]) were imaged on a Zeiss LSM 700 inverted confocal microscope using a 20×/0.8NA Plan-APOCHROMAT objective lens with refractive index correction. 2μm z-stacks were acquired through the entire thickness of the section at 1AU (0.313×0.313×2μm). Three brain sections, separated by at least 100μm, were imaged, cell profiles counted, and averaged per animal. Manual counts were performed using the ‘cell counter’ plugin in FIJI software version 2.9.0 (https://fiji.sc/, RRID:SCR_002285). Images were cropped by layer based on CTIP2 immunostaining and DAPI, to perform layer-specific analysis. The width of the cortical crop was held constant (321µm), while the thickness varied to capture the full depth of each layer. The total number of DAPI nuclei, FOS^+^ cells and GFP^+^ cells, as well as FOS^+^; GFP^+^ double-labeled cells, were counted. The marker of interest was considered a positive count only if there was both immunofluorescence signal and a DAPI^+^ nucleus.

### Statistical Analyses

Data are reported as meanL±Lstandard error to the second decimal place. Sample sizes are reported in the figures. An individual animal represents a single sample. Sample sizes were determined using a power analysis at α=0.05 and 1-β=0.8 (SPH Analytics, statistical power calculator using average values). For all analyses, test statistics are reported to the second decimal place and *p* values are reported to the fourth decimal place. Statistical analyses were performed using GraphPad Prism software version 9.1.2 (GraphPad Software, Inc, La Jolla, CA). For each statistical analysis, a D’Agostino & Pearson normality test (n≥8) or a Shapiro-Wilk normality test (n<8), was performed. One-tailed Mann-Whitney (non-parametric; test statistic: *U*), Kruskal-Wallis (non-parametric; test statistic: *H*), followed by a post hoc Dunn’s multiple comparisons test if omnibus test detected a significant difference, two-tailed unpaired *t*-tests (parametric; test statistic: *t*), ordinary one-way ANOVA (parametric; test statistic: *F*), followed by a post hoc Tukey multiple comparisons tests if omnibus test detected a significant difference, Kolmogorov-Smirnov (non-parametric; test statistic *D*), and two-way ANOVA, followed by Šídák’s multiple comparisons test, were used.

## RESULTS

### Developmental Trajectories of Contextual Fear Memory Persistence and Remote Memory

To determine the precise developmental onset of contextual fear memory persistence capabilities in WT mice, Shock mice at different developmental ages were conditioned and tested 7-d later. Freezing responses on testing day were compared to age-matched No-Shock mice. Contextual fear memory persistence capabilities were considered present when the Shock group exhibited significantly increased freezing responses compared to the No-Shock group. There was no significant difference in percentage time freezing between the Shock (1.65 ± 1.05) and No-Shock (0.56 ± 0.34) groups 7-d after training on P15 (*U* = 26.00; *p* = 0.4168; Fig. 1B), contrasting with the significant differences observed in mice trained on P35 (*U* = 1.00; *p* < 0.0001; No-Shock: 0.44 ± 0.16; Shock: 16.92 ± 4.80; Fig. 1C). These results demonstrate that the ability to retain a contextual fear memory for at least 7-d emerges between P15-P35 in C57BL/6J mice, consistent with Akers et al., using a mixed background strain [20].

We next trained separate cohorts of mice at various ages between P15-P35, testing for persistent memory. For mice trained on P20, the Shock group (2.80 ± 1.56) exhibited no significant difference in percentage time freezing compared to the No-Shock group (0.68 ± 0.14; *U* = 22.00; *p* = 0.4242; Fig. 1D). However, mice trained on P15 or P20 and tested 1-d later exhibited contextual fear memory formation capabilities (Fig. S1A-B), demonstrating that the lack of persistent memory expression at these ages is not due to memory formation deficits. Remarkably, when mice were trained 1-d later (P21), a significant difference between the two groups was observed during persistent memory testing (*U* = 0; *p* < 0.0001; No-Shock: 0.09 ± 0.05; Shock: 23.08 ± 6.87; Fig. 1E). Notably, on training day there were no differences in spontaneous baseline freezing during the first two minutes of chamber exposure (*U* = 71.50; *p* = 0.3571; No Shock: 1.39 ± 0.58; Shock: 2.17 ± 1.28; Fig. S1C). As this represents the time prior to shock administration, the freezing occurring in the Shock group on testing day is a learned response rather than spontaneous freezing. Mice trained at older ages similarly exhibited little to no baseline freezing (data not shown). To determine whether there was a developmental change in the robustness of contextual fear memory expression following the initial onset of persistence, we next compared age, training condition (Shock or No Shock), and interaction effects of the percentage time freezing of mice trained on P21, P22, P23, or P35 and tested 7-d later. Two-way ANOVA revealed a main effect of training day condition (*p* < 0.0001), but no age (*p* = 0.4063) or interaction effect (*p* = 0.4758), indicating no further maturation of persistent memory expression over the two weeks following onset (P21 No-Shock: 0.09 ± 0.05; P21 Shock: 23.08 ± 6.87; P22 No-Shock: 1.43 ± 0.98; P22 Shock: 29.08 ± 7.43; P23 No-Shock: 0.46 ± 0.32; P23 Shock: 31.97 ± 6.94; P35 No-Shock: 0.44 ± 0.16; P35 Shock: 16.92 ± 4.80; Fig. 1F). Together, these data show surprisingly rapid onset of contextual fear memory persistence capabilities at P21, with comparable memory expression as older ages.

Remote contextual fear memory capabilities are present at P25 but not at P21 in WT mice using a 5-shock paradigm [20]. We determined whether remote fear memory is present within this timeframe using the 3-shock paradigm. Thirty days following training at P23, one of the earliest ages that contextual fear memory persistent capabilities are present, there was a significant increase in percentage time freezing in Shock (7.05 ± 2.24) compared to No-Shock (0.02 ± 0.02; *U* = 28; *p* < 0.0001; Fig. 2A) mice. Because the freezing response appeared blunted compared to that observed after the 7-d training-testing interval (Fig. 1G), we measured the freezing response of P23 and P35 Shock mice at different training-testing intervals (7-, 14-, 30-d) to determine if there were any developmental differences between these two ages in fear memory expression of longer-term memory (Fig. 2B). Two-way ANOVA revealed no main effect of training-testing interval (*p* = 0.0567) or age (0.4073), but there was an interaction effect (*p* = 0.0084). Post-hoc multiple comparisons showed that mice trained at P23 exhibited less freezing (*p* = 0.0004) if tested 30-d after training (7.05 ± 2.24) compared to mice tested 7-d after training (31.97 ± 6.94). Thus, P23 mice exhibit blunted remote fear memory expression compared to memory persistence, while 14-d memory retention resulted in intermediate fear expression (15.34 ± 4.26). In contrast, post-hoc multiple comparisons revealed no significant differences when mice were trained on P35 and tested 7-, 14-, or 30-d later (7-d: 16.92 ± 4.80; 14-d: 28.92 ± 6.78; 30-d: 18.98 ± 5.08), demonstrating that at P35, memory expression is as robust for remote memory as for shorter intervals of memory expression.

**Fig. 2:**
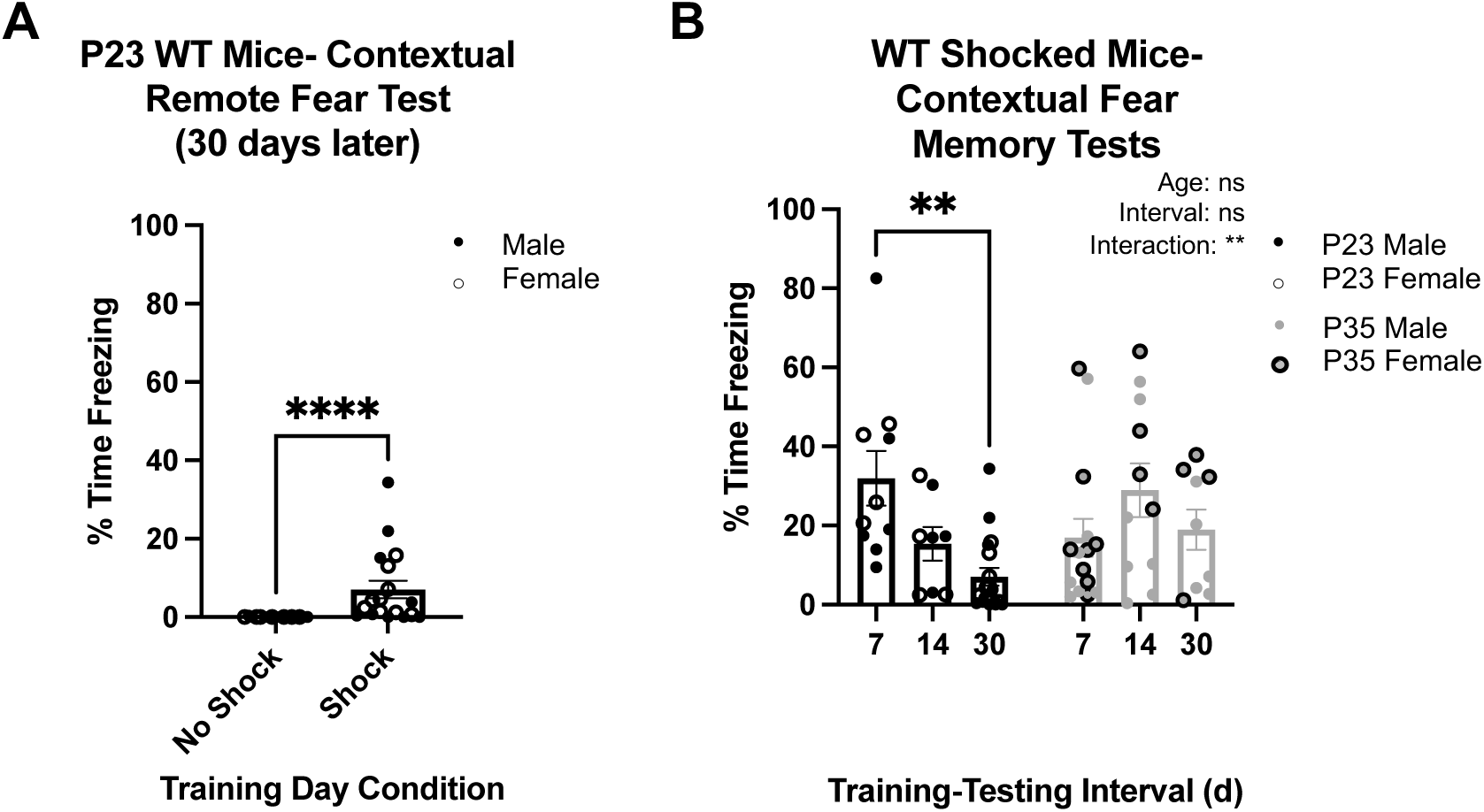
Traces of remote contextual fear memory are present at P23 but is still developing compared to P35. **A** Quantification of the percentage time freezing on testing day of No Shock (n = 17) and Shock (n = 18) mice trained on P23 and tested 30 d later. The Shock group exhibited significantly more freezing than No Shock (\*\*\*\**p* < .0001). **B** Quantification of the percentage time freezing on testing day of Shock mice conditioned on P23 or P35 and tested 7 (n = 10), 14 (n = 8), or 30 (n = 18) d later. There is a significant difference between 1 and 30 d training intervals at P23, and no differences were determined at P35 (\*\**p* < .01, ns, not significant).

### Sustained *Met* Expression Does Not Disrupt Developmental Trajectory of Expressed Fear Memory Retention

MET protein in the cerebral cortex downregulates starting by P21 and is greatly reduced by P35 [22]. With the developmental trajectories for persistent and remote memories defined and coinciding temporally with MET downregulation and given the effects of sustained MET expression on disrupting critical period timing following monocular deprivation [25], we tested the hypothesis that normal downregulation of MET expression in the cortex is necessary for emergence of contextual fear memory persistence and remote memory during development. Separate cohorts of cto-*Met* (sustained MET expression) and littermate control mice were trained between P23-P90 and tested 7- or 30-d later. Because there were no genotype differences found at any age in the No-Shock mice (data not shown), only genotype effects between age-matched Shock mice were compared. There was no difference between control (32.75 ± 4.19) and cto-*Met* (32.39 ± 7.10) Shock mice in percentage time freezing for those trained on P23, one of the earliest ages that memory persistence is present in WT mice, and tested 7-d later (*t* = 0.05; *p* = 0.9640; Fig. 3A). Next, P35 mice, an age at which there is robust remote memory expression in WT mice (Fig. 2C), were tested for remote memory. Remote memory was intact, with no difference between control (23.69 ± 5.97) and cto-*Met* (21.01 ± 5.59) Shock mice in percentage time freezing on testing day (*t* = 0.33; *p* = 0.7480; Fig. 3B). Together, these data indicate that downregulation of MET expression is not required for emergence of contextual fear memory persistence or remote memory capabilities. Lastly, we determined there was no difference between P90 control (18.46 ± 4.06) and cto-*Met* (14.20 ± 3.65) Shock mice in percentage time freezing on testing day 7-d later *(t* = 0.78; *p* = 0.4448; Fig. 3C), indicating downregulation of MET is not necessary for memory persistence capabilities in adulthood.

**Fig. 3:**
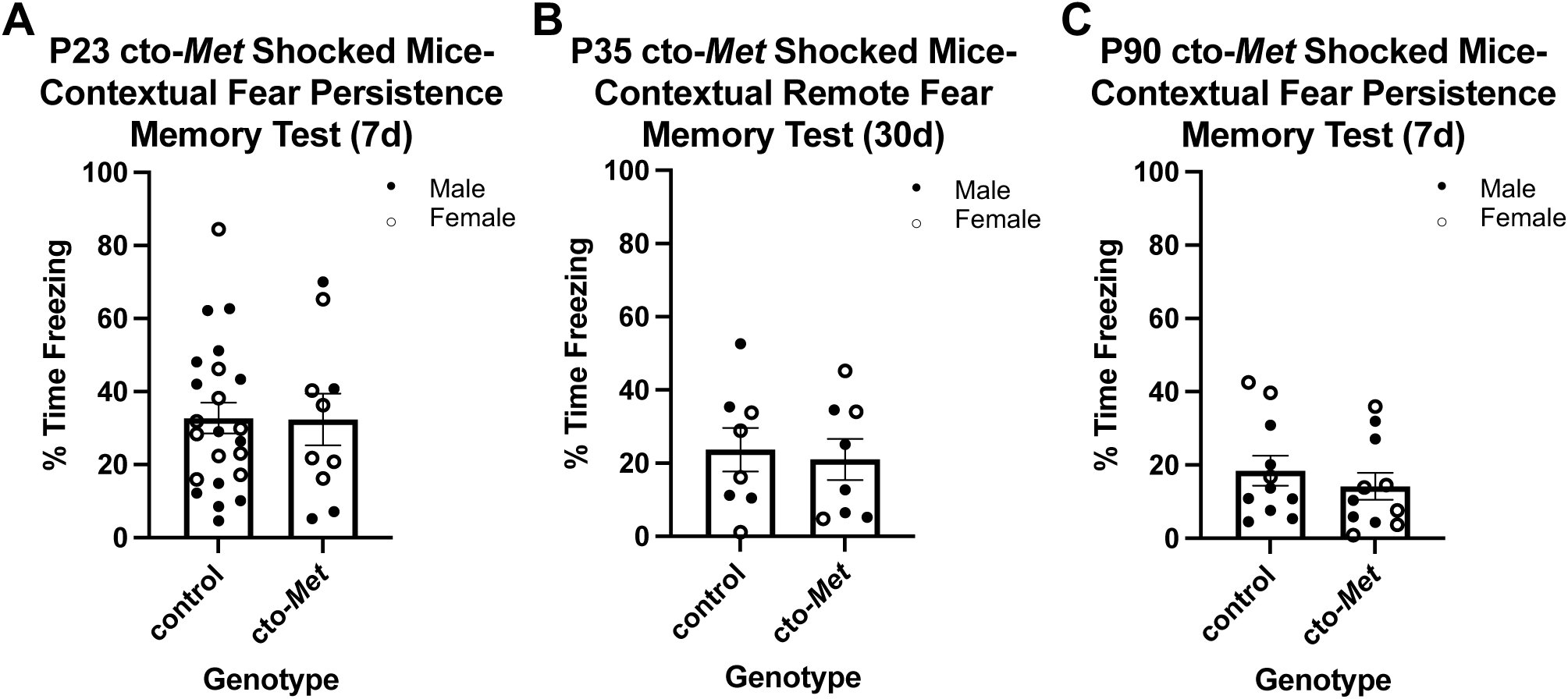
Sustaining MET past its normal temporal peak in the cortex does not affect contextual fear memory persistence developmentally or in adulthood. **A** Quantification of the percentage time freezing on testing day of control (n = 23) and cto-*Met* (n = 10) genotypes of Shock mice conditioned on P23 and tested 7 d later. No significant difference between genotypes was determined. **B** Quantification of the percentage time freezing on testing day of control (n = 8) and cto-*Met* (n = 8) genotypes of Shock mice conditioned on P35 and tested 30 d later. No significant difference between genotypes was determined. **C** Quantification of the percentage time freeing on testing day of control (n = 11) and cto-*Met* (n = 11) genotypes of Shock mice conditioned on P90 and tested 7 d later. No significant difference between genotype was determined.

### *Met* Deletion Results in Delayed Disruption of Fear Memory Expression

We next probed the timing of when the previously reported adult contextual fear deficits arise in the *Nestin*-cre; *Met*^fx^ line [28-29]. Fear memory deficits have been reported in adult male *Nestin*-cre mice [33], but the impact of the *Nestin*-cre driver during development and in adult females had not been determined. First, we replicated the contextual fear memory deficits previously reported in adult male *Nestin*-cre mice [33], demonstrating a significant effect of the Cre genotype on percent freezing on testing day, one day after training on P90 (*D* = 0.86; *p* = 0.0152; Cre-: 28.36 ± 7.06; Cre+: 4.71 ± 2.62; Fig. S2A). There was, however, a sexually dimorphic effect of Cre at this age, such that the *Nestin*-cre driver exhibited normal contextual fear memory formation in female mice (*t* = 1.49; *p* = 0.1506; Cre-: 16.87 ± 4.13; Cre+: 10.05 ± 2.21; Fig. 4A). Further, neither male nor female *Nestin*-cre driver mice exhibited a deficit in fear memory formation during development. Specifically, at P23, Cre- and Cre+ mice exhibited no difference in contextual fear memory formation (*t* = 0.32; *p* = 0.7501; Cre-: 31.80 ± 9.24; Cre+: 29.07 ± 3.89; Fig. S2B) or persistence (*D* = 0.42; *p* = 0.4680; Cre-: 32.60 ± 10.23; Cre+: 21.48 ± 6.20; Fig. S2C). Similarly, there was no significant effect of the *Nestin*-cre driver on remote contextual fear memory at P35 (*D* = 0.38; *p* = 0.5077; Cre-: 10.67 ± 2.39; Cre+: 17.21 ± 8.00; Fig. S2D) and P50 (*D* = .34; *p* = 0.6277; Cre-: 34.02 ± 8.79; Cre+: 27.24 ± 7.31; Fig. S2E). Together, these data demonstrate the utility of the *Nestin*-cre driver line to conditionally delete *Met* developmentally in both sexes, and in adult females to study the effects on contextual fear memory.

**Fig. 4:**
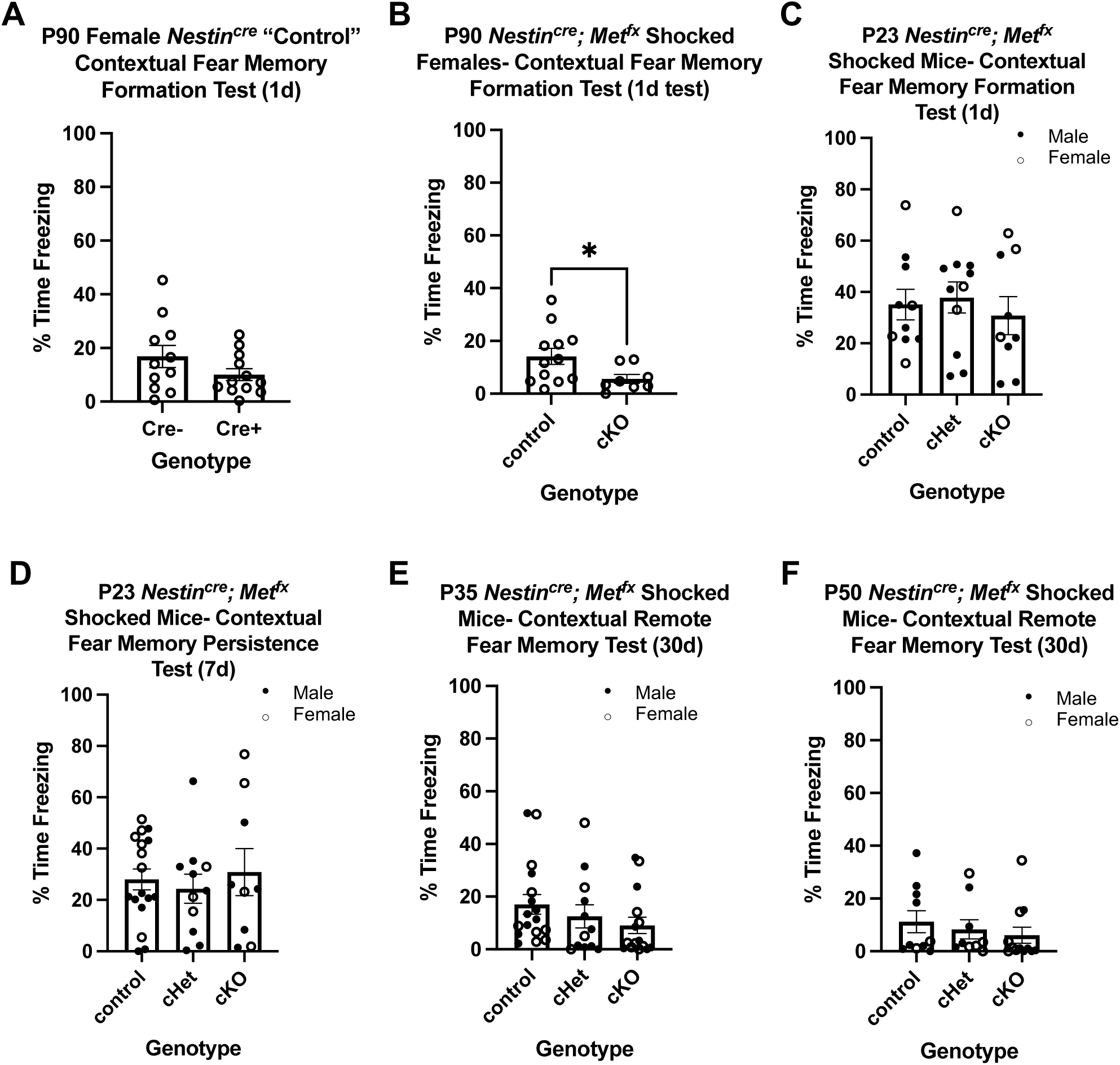
Absence of MET expression in neurons impacts contextual fear memory in adult females but not in developing male and female mice. **A** Quantification of the percentage time freezing on testing day of Cre- (n = 11) and Cre+ (n = 12) *Nestin^cre^*female Shock mice conditioned on P90 and tested 1 d later. No significant difference between genotypes was determined (*p* = .1506). **B** Quantification of the percentage time freezing on testing day of control (n = 12) and cKO (n = 8) *Nestin^cre^*/*Met^fx^*female Shock mice conditioned on P90 and tested 1 d later. cKO exhibited significantly less freezing than control (*p* = 0.0455). **C** Quantification of the percentage time freezing on testing day of control (n = 10), cHet (n = 11), and cKO (n = 9) *Nestin^cre^*/*Met^fx^* Shock mice conditioned on P23 and tested 1 d later. No significant difference across genotypes was determined (*p* = .7440). **D** Quantification of the percentage time freezing on testing day of control (n = 17), cHet (n = 11), and cKO (n = 9) *Nestin^cre^*/*Met^fx^* Shock mice conditioned on P23 and tested 7 d later. No significant difference across genotypes was determined (*p* = .7734). **E** Quantification of the percentage time freezing on testing day of control (n = 17), cHet (n = 12), and cKO (n = 15) *Nestin^cre^*/*Met^fx^* Shock mice conditioned on P35 and tested 30 d later. No significant difference across genotypes was determined (*p* = .0989). **F** Quantification of the percentage time freezing on testing day of control (n = 10), cHet (n = 9), and cKO (n = 12) *Nestin^cre^*/*Met^fx^* Shock mice conditioned on P50 and tested 30 d later. No significant difference across genotypes was determined (*p* = .2286).

In P90 adult female mice, there are deficits in contextual fear memory formation when *Met* is conditionally deleted embryonically (*t* = 2.15; *p* = 0.0455; control: 14.16 ± 3.01; cKO: 5.66 ± 1.67; Fig. 4B). Notably, on training day control and cKO mice displayed no baseline freezing responses (0%; Fig. S2F), demonstrating freezing on testing day is a learned response. In contrast, following training on P23, there was no effect of genotype on percentage time freezing 1-d (*F* = 0.30; *p* = 0.7440; control: 35.10 ± 5.9414; cHet: 37.82 ± 6.05; cKO: 30.78 ± 7.39; Fig. 4C) and 7-d (*F* = 0.26; *p* = 0.7734; control: 28.00 ± 4.07; cHet: 24.37 ± 5.64; cKO: 30.87 ± 9.14; Fig. 4D). Similarly, there was no effect of genotype on percentage time freezing in mice tested for remote memory 30-d after training on P35 *(H* = 4.63; *p* = 0.0989; control: 17.05 ± 3.77; cHet: 12.51 ± 4.40; cKO: 9.07 ± 3.14; Fig. 4E) or ∼P50 (post-pubertal adolescence [38]; *H* = 2.95; *p* = 0.2286; control: 11.23 ± 4.18; cHet: 8.31 ± 3.64; 6.09 ± 3.05; Fig. 4F). These data demonstrate a substantial delay in females in contextual fear memory deficits, first acquired in young adults, following embryonic elimination of *Met* in neural cells.

### Differences Between Developmental and Adult Infragranular mPFC in Activation of Total Neurons and *Met*^GFP^ Subtype During Fear Memory Expression

mPFC contributes to contextual fear memory circuitry [39-40], and MET is enriched in subcerebral projection neurons in infragranular layers of mPFC throughout postnatal development in mice [41]. We hypothesized that age-dependent differences in the MET population could contribute to the observed female adult-specific deficit in fear memory. We first determined if there were differences in the percentage of MET-expressing neurons at P35 compared to adult mice. Analyses using the *Met*^GFP^ line to visualize GFP in *Met*-expressing cells (Fig. 5A) revealed no significant difference in MET-GFP cell density in layer 5 (P35: 15.59 ± 0.58; P90: 17.61 ± 1.13; *t* = 1.50; *p* = 0.1691; Fig. 5B) or layer 6 (P35: 14.04 ± 1.00; P90: 8.85 ± 1.78; *D* = 0.67; *p* = 0.1429; Fig. 5C).

**Fig. 5:**
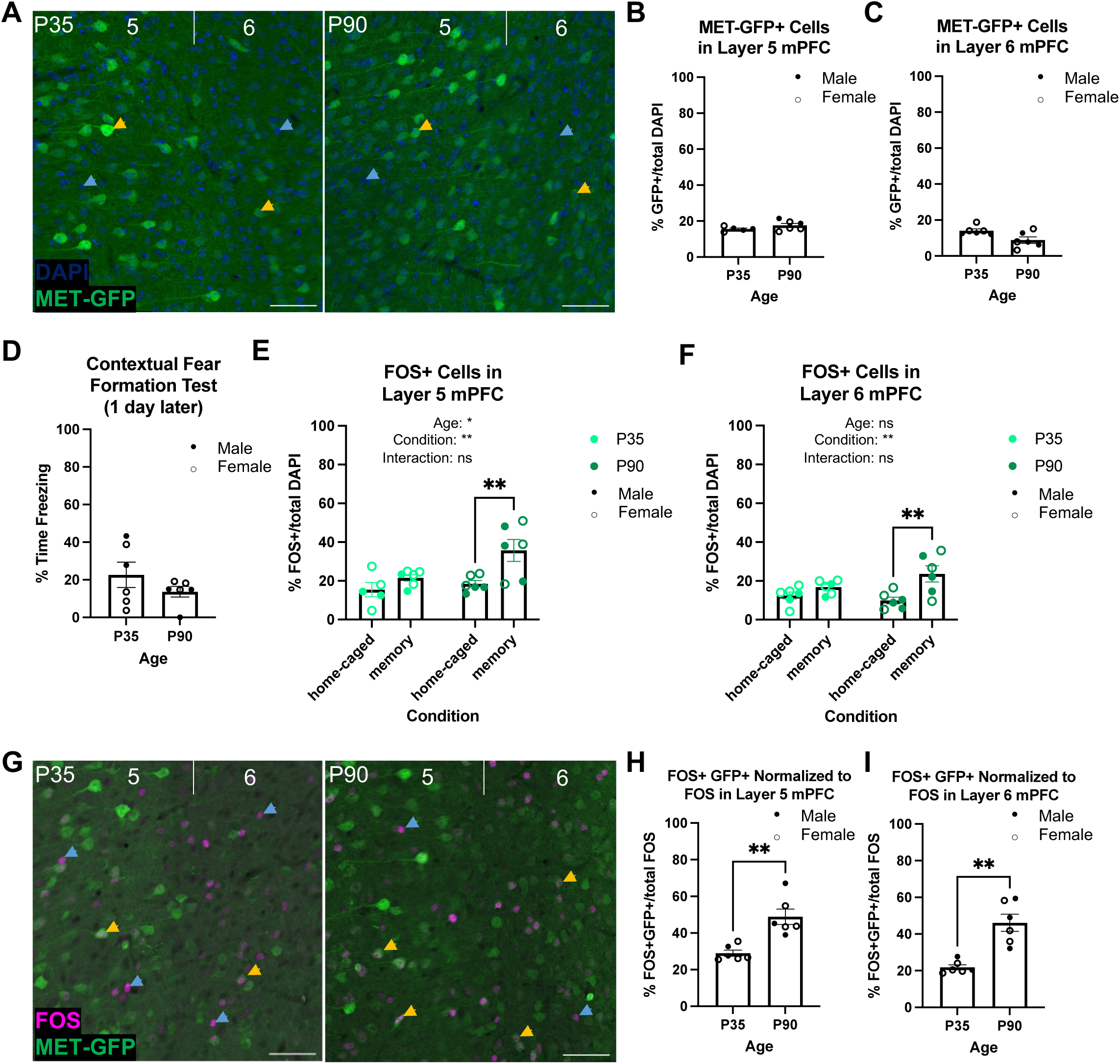
MET-GFP and FOS expression in infragranular layers 5 and 6 of mPFC at P35 compared to P90. **A** Exemplary images of DAPI (blue) and MET-GFP (green) in layers 5 and 6 mPFC at P35 (left) and P90 (right). Blue arrows denote DAPI nuclei, yellow arrows denote GFP^+^ cells, scale bars = 50 µm. **B** Quantification of the percentage of MET-GFP^+^ cells in layer 5 mPFC at P35 (n = 5) and P90 (n = 6). No significant difference between ages was determined. **C** Quantification of the percentage of MET-GFP^+^ cells in layer 6 mPFC at P35 (n = 6) and P90 (n = 6). No significant difference between ages was determined. **D** Quantification of the percentage time freezing of *Met*^GFP^ Shock mice conditioned on P35 (n = 6) or P90 (n = 6) and tested 1 d later. No significant difference between ages was determined. **E** Quantification of the percentage of FOS^+^ cells in layer 5 mPFC in P35 and P90 home-caged and contextual fear memory formation tested mice (P35 home-caged: n = 5; P35 memory: n = 6; P90 home-caged: n = 6; P90 memory: n = 6). There were significant age and condition effects, with a significant difference between P90 home-caged and P90 memory tested mice (\**p* < .05, \*\**p* < .01, ns, not significant). **F** Quantification of the percentage of FOS^+^ cells in layer 6 mPFC in P35 and P90 home-caged and contextual fear memory formation tested mice (P35 home-caged: n = 6; P35 memory: n = 6; P90 home-caged: n = 6; P90 memory: n = 6). There was a significant condition effect, with a significant difference between P90 home-caged and P90 memory tested mice (\*\**p* < .01, ns, not significant). **G** Exemplary images of FOS (magenta) and MET-GFP (green) in layers 5 and 6 mPFC at P35 (left) and P90 (right). Blue arrows denote FOS^+^ cells, yellow arrows denote FOS^+^;GFP^+^ colocalized cells, scale bars = 50 µm. **H** Quantification of the percentage of FOS^+^;GFP^+^ colocalized cells out of total FOS-expressing cells in layer 5 mPFC in mice conditioned on P35 (n = 6) or P90 (n = 6) and tested 1 d later and in layer 6 mPFC (**I**). There was a significant difference between ages in both layers (\*\**p* < .01).

We next probed whether there were age differences in the percentage of MET-expressing cells that are activated during memory expression. *Met*^GFP^ mice were conditioned on P35 or P90, returned to the chamber for memory testing 1-d later and sacrificed 90 minutes after chamber exposure on testing day. Memory-induced FOS^+^ cells were quantified and compared to baseline levels of FOS^+^ cells in age-matched mice that remained in their home cage, without chamber exposure or conditioning during this time. We performed an initial assessment, revealing that the minor handling of the mice performed during the testing day did not, on its own, result in obvious changes in the number of FOS+ cells in the mPFC (data not shown). Importantly, there was no significant difference in freezing responses of conditioned mice during 1-d memory testing between P35 (22.63 ± 6.72) and P90 (13.64 ± 2.84), demonstrating that fear expression behavior is comparable at both ages (*D* = 0.50; *p* = 0.4740; Fig. 5D). We first considered the FOS-expressing population independent of GFP expression. Two-way ANOVA revealed a main effect of age (*p* = 0.0268) and condition (memory-tested versus homecage; *p* = 0.0043), but no interaction effect (*p* = 0.1391) on the percentage of FOS^+^ cells in layer 5 (Fig. 5E). Unexpectedly, post-hoc analyses revealed no significant difference in percentage of FOS^+^ cells between memory-tested (21.51 ± 1.68) and home-caged (15.40 ± 3.73; *p* = 0.4453) mice at P35, contrasting with the significantly higher percentage of FOS^+^ cells in memory-tested (35.69 ± 5.66; *p* = 0.0051) compared to home-caged (18.48 ± 1.63) mice at P90. In layer 6 mPFC, two-way ANOVA revealed a main effect of condition (*p* = 0.0020), but no age effect (*p* = 0.4196) or interaction (*p* = 0.0812; Fig. 5F). Again, post-hoc analyses revealed no significant difference in percentage of FOS cells between home-caged (12.44 ± 1.89) and memory (16.81 ± 1.44; *p* = 0.4223) mice at P35, in contrast to a significantly higher percentage of FOS^+^ cells in memory-tested (23.59 ± 4.17; *p* = 0.0022) compared to home-caged (9.86 ± 1.73) mice at P90. Together, these results indicate that cells in infragranular mPFC are preferentially engaged during memory expression at P90, but not P35, compared to baseline. Finally, we quantified the percentage of FOS^+^;GFP^+^ double-labeled cells in mPFC at P35 and P90 during 1-d memory expression (Fig. 5G). There was a significant increase in FOS^+^; GFP^+^ cells, normalized to total FOS^+^ cells, at P90 compared to P35 in layer 5 (P35: 28.98 ± 1.67, P90: 48.85 ± 4.23; *t* = 4.37 *p* = 0.0014; Fig. 5H) and layer 6 (P35: 21.84 ± 1.38, P90: 46.11 ± 4.66; *D* = 1.00; *p* = 0.0022; Fig. I). These data indicate that MET-GFP^+^ cells in mPFC are preferentially recruited during 1-d fear memory expression in adults, but not at P35.

## DISCUSSION

The present study provides a new understanding of the temporal profile for the acquisition of memory capabilities in developing mice and the role of the receptor tyrosine kinase, MET, in temporally distinct mediation of fear memory expression. The abrupt onset of memory persistence between P20-P21 occurs after mice can retain a memory for 1-d but prior to having the capacity for fully expressed remote memory. This is independent of weaning, as shifting weaning a day earlier or later had no impact on the onset of contextual fear memory persistence (data not shown). There were also no trending sex effects for this developmental onset using the current paradigm, with males and females thus analyzed together. The lack of sex differences could, in part, be due to the fact that this onset occurs prior to puberty. However, we did not have the statistical power to test for small effect size sex differences. Interestingly, in contrast to the 1-d transition for onset of memory persistence, remote memory becomes fully expressed gradually over the days following initial onset. Together, these data indicate that longer-term memory capabilities develop in a stepwise fashion, first with emergence of shorter-term abilities that mature into longer-term capabilities over time. The current findings also provide a necessary temporal framework for future studies probing the mechanisms that underlie these transitions and for determining biological and environmental factors that accelerate or delay this trajectory. For example, rapid maturation in circuitry that connects the mPFC, hippocampus, and basolateral amygdala could be one possible underlying change that occurs between P20 and P21, enabling longer-term memory capabilities. There also may be molecular changes that underlie expression of memory persistence. Interestingly, the onset of fear memory precision capabilities, in which mice display significant freezing behavior only in the chamber where the shocks were administered, compared to having generalized fear in a different chamber environment, is not present at P20, but emerges by P24 and correlated with hippocampal maturation [42]. Therefore, developmental changes in hippocampus during this time period may also be involved in the initial onset of contextual fear memory persistence. Together, the data identify a potential sensitive period in memory development, during which disruptions in the maturational processes that allow this cognitive function to become fully expressed would have a profound impact on learning and memory capabilities.

Based, in part, on the correlation between the timing of MET expression downregulation in the cortex and the onset of contextual fear memory persistence, we focused on a potential role for MET in the development of the cognitive capacity of fear memory persistence. Contrary to our hypothesis, based on our studies on critical periods in visual cortex [25], an “off” signal of *Met* is not necessary for normal contextual fear memory during development or into adulthood. However, we did not test for remote fear memory deficits at P90 or perform any conditioning beyond the age of P90. Thus, additional studies will be needed to determine whether sustained *Met* signaling impacts contextual fear memory later in adulthood. Further, while *Met* expression is necessary for contextual fear memory expression in young adult female mice, embryonic deletion or reduction of *Met* had no impact on contextual fear memory expression through the late adolescent period in male and female mice. Given that the Nestin-Cre driver line exhibits a sexually dimorphic impact on freezing, but only in adult mice, whether the deficit delay from *Met* deletion is sex-specific or occurs in both males and females will be determined in future studies with a different driver line that does not exhibit sex-specific deficits. The temporal delay of female deficits is similar to a study in which behavioral deficits were observed in adult, but not juvenile, mice that are haploinsufficient in *Myt1l*, a gene highly expressed early postnatally but not in adults [43]. For both MET and MYT1L, reduction in normal protein expression leads to the emergence of behavioral deficits only after circuit maturation is complete, well beyond the period of normal peak protein expression.

The finding of increased activation of infragranular mPFC neurons during 1-d memory expression compared to baseline at P90, but not P35, support a greater reliance on engaging mPFC circuitry for memory expression in adults compared to the early adolescent period. We note, however, that we compared conditioned mice to home-cage controls at both ages, and thus cannot disentangle whether increased mPFC activation at P90 is due to context memory alone, which is one aspect in contextual fear memory processing, or due specifically to the fear memory recall. Future studies will determine in which aspect of fear memory encoding these mPFC neurons are involved. Regardless, these results demonstrate a previously unknown change in male and female mice that occurs between P35 and P90 in the engagement of mPFC neuron activation during contextual fear memory recall. Because male and female data did not show any differences between sexes (data were collapsed), this suggests that a similar increased mPFC engagement occurs in males and females by P90. Consistent with our results, inactivation of mPFC after conditioning in post-weanling rats does not affect 1-d memory expression [44]. We suggest that adolescence represents a second sensitive period, during which fear memory capabilities are present, but the underlying brain circuits recruited are different than in adults. MET signaling participates in molecular maturation of synaptic elements, regulating β-catenin and N-cadherin interactions, trafficking of glutamate receptor subunits and expression of small GTPases [27, 45-46]. If, during this sensitive period, circuits subserving adult fear memory do not develop properly, for example due to deletion of *Met* expression, deficits may arise as contextual fear memory becomes more dependent upon mPFC involvement. Major pruning in mPFC projections to the basal amygdala, connectivity that is involved in adult fear memory circuitry, occurs between P45-P90 [47]. This period of refinement represents a time during which circuitry is vulnerable and, if disrupted, could lead to adult-specific deficits.

*MET* was initially considered an autism spectrum disorder (ASD) risk gene and exhibits reduced expression in postmortem tissues from temporal neocortex in ASD and Rett Syndrome [48-51], but it more likely serves as an important receptor for modulating the maturation of subsets of synapses in the developing cerebral cortex, leading to circuit development vulnerabilities if signaling is disrupted. Studies of MET function in the cerebral cortex therefore has been studied largely in a developmental context. The adult-onset deficits in fear memory formation observed in female *Met* null mice could, however, reflect a previously unrecognized adult-specific function of MET. Indeed, the findings that MET expression is enriched in the FOS^+^ cells in infragranular mPFC during the expression of fear memory in adults compared to adolescents, even though the percentage of MET-GFP expressing cells between ages is the same, indicate an adult-specific engagement of MET-expressing cells in mPFC during fear memory expression. Because cortical neurons with sustained MET expression have more biochemically immature synaptic properties [24], adult expression of MET may maintain subsets of synapses in a more plastic state during adult learning and memory. Overall, these results do not model the clinical presentations of ASD, but rather model the development of circuit vulnerability involving typical learning and memory capabilities.

The present study advances insight into the precise timing of the ontology of contextual fear memory capabilities, facilitating the design of additional studies to determine the molecular mechanisms involved in the onset and continued expression of this cognitive function into adulthood. Additionally, the finding that not all functional deficits observed in adults are expressed during development emphasizes the need for careful temporal analyses of the timing onset of functional disturbances. Experimental data that precisely define typical development trajectories of additional cognitive abilities would further benefit studies aiming to address mechanisms underlying neurodevelopmental disorders and the cognitive deficits that are often associated with them. This study focused on contextual fear memory, and thus, does not exclude MET from having roles in regulating the onset of other cognitive functions and the timing of distinct critical periods. We also note that while the present study focused on mPFC, other brain regions may also contribute to the observed female adult deficits when *Met* is conditionally deleted. Future studies will address the impact of modulating neuronal activity of neuronal subtypes expressing MET in mPFC on contextual fear memory in adults, as well as on other cognitive assays across development.

## ACKNOWLEDGEMENTS

We thank Amanda Whipple for assistance with animal husbandry and genotyping. We thank Dieter Hertling for technical assistance with the contextual fear memory assay. We thank Dr. Allison Knoll and Miranda Villanueva for technical assistance in setting up the contextual fear memory chambers and paradigms. Lastly, we thank Dr. Esteban Fernandez of the Cellular Imaging Core at The Saban Research Institute (TSRI), Children’s Hospital Los Angeles for assistance with image acquisition and analysis.

## AUTHOR CONTRIBUTIONS

All authors contributed to conceiving and designing the research. A.L.L. performed the experiments and analyses. All authors interpreted the data. A.L.L. drafted the manuscript. All authors revised the manuscript for important intellectual content. All authors approved the final version to be published. All authors agree to be accountable for all aspects of the work.

