## Supplemental Figure 1 for "Developmental and molecular contributions to contextual fear memory emergence in mice"

### Slide 1
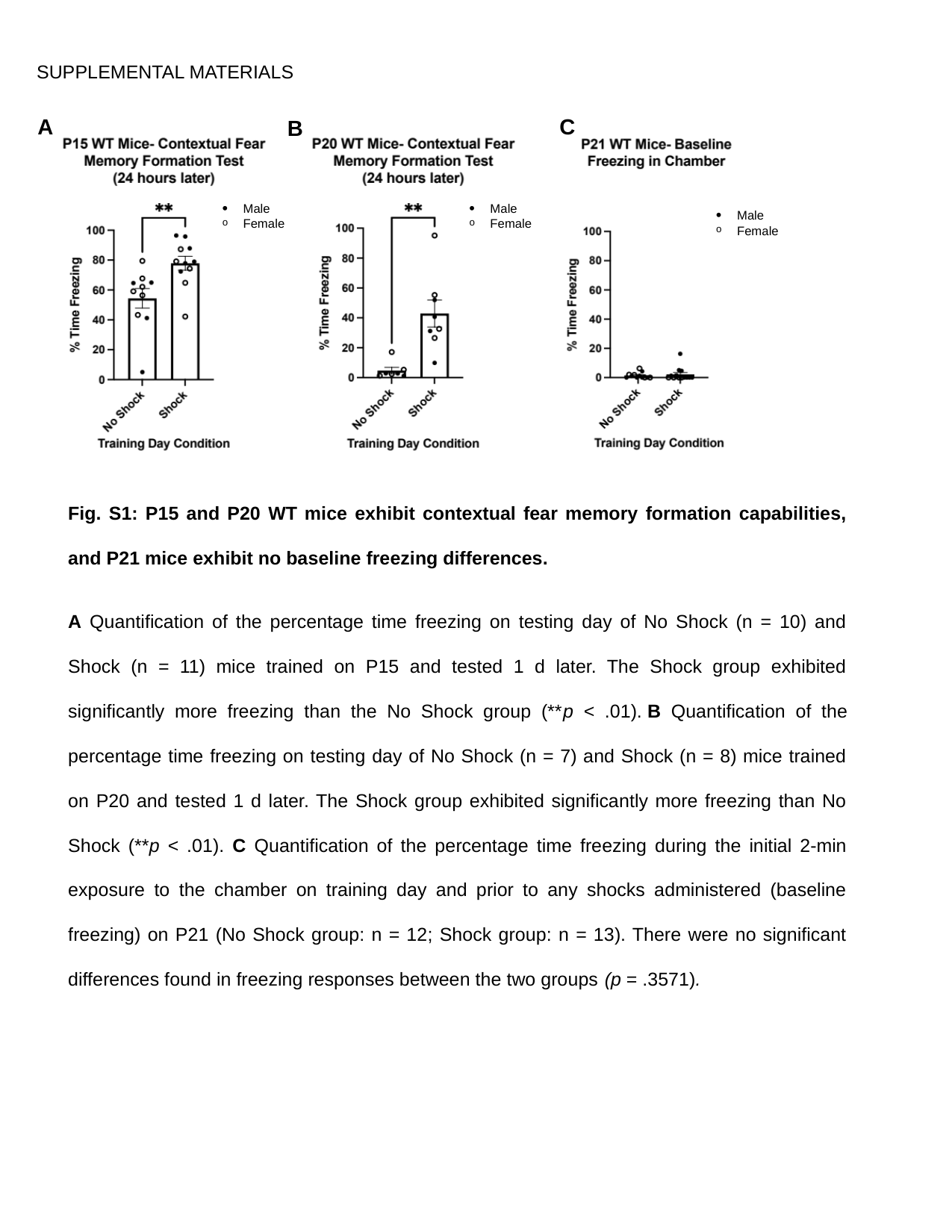

SUPPLEMENTAL MATERIALS
A
C
B
Male
Female
Male
Female
Male
Female
Fig. S1: P15 and P20 WT mice exhibit contextual fear memory formation capabilities, and P21 mice exhibit no baseline freezing differences.
A Quantification of the percentage time freezing on testing day of No Shock (n = 10) and Shock (n = 11) mice trained on P15 and tested 1 d later. The Shock group exhibited significantly more freezing than the No Shock group (**p < .01). B Quantification of the percentage time freezing on testing day of No Shock (n = 7) and Shock (n = 8) mice trained on P20 and tested 1 d later. The Shock group exhibited significantly more freezing than No Shock (**p < .01). C Quantification of the percentage time freezing during the initial 2-min exposure to the chamber on training day and prior to any shocks administered (baseline freezing) on P21 (No Shock group: n = 12; Shock group: n = 13). There were no significant differences found in freezing responses between the two groups (p = .3571).
