## Supplemental Figure 2 for "Developmental and molecular contributions to contextual fear memory emergence in mice"

### Slide 1
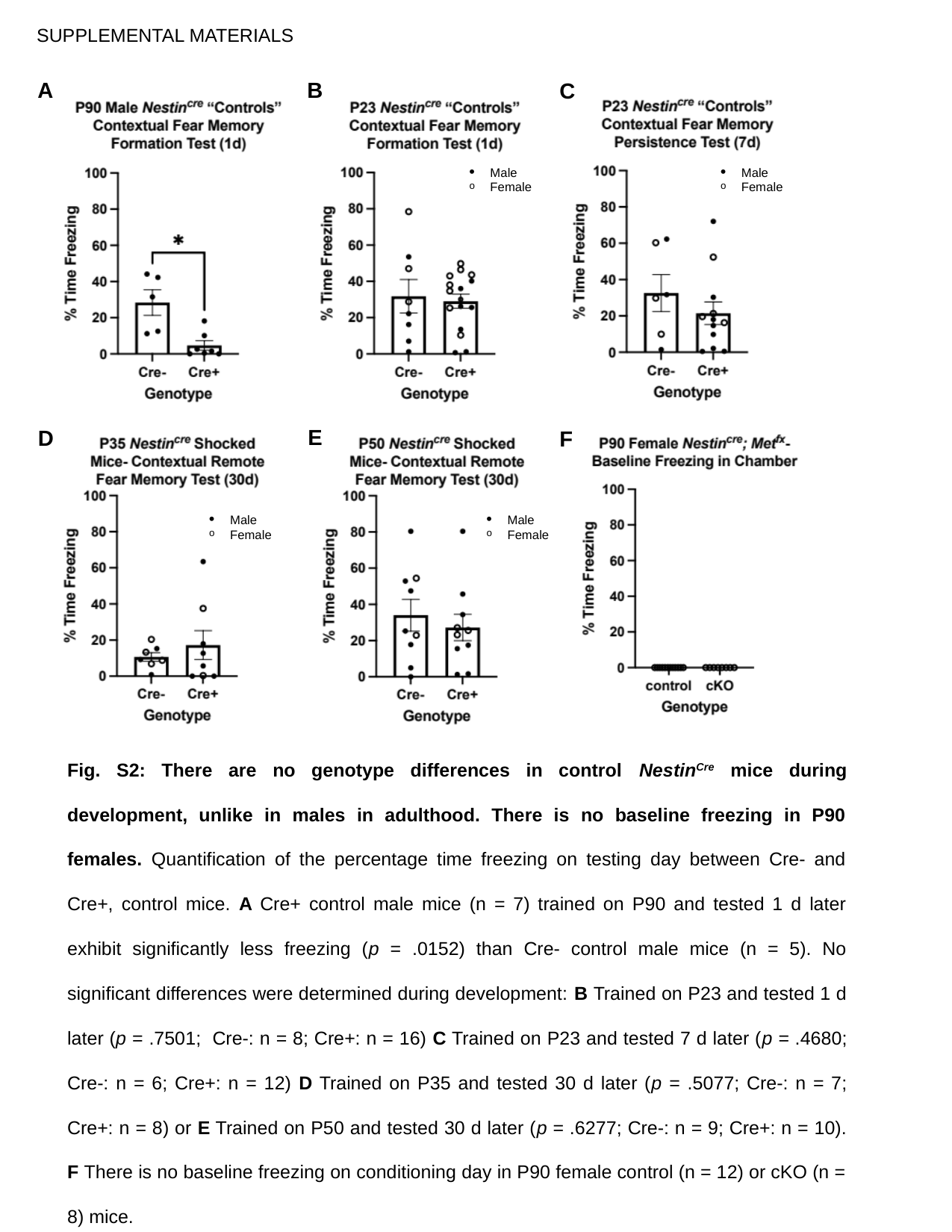

SUPPLEMENTAL MATERIALS
A
B
C
Male
Female
Male
Female
E
D
F
Male
Female
Male
Female
Fig. S2: There are no genotype differences in control Nestin^Cre^ mice during development, unlike in males in adulthood. There is no baseline freezing in P90 females. Quantification of the percentage time freezing on testing day between Cre- and Cre+, control mice. A Cre+ control male mice (n = 7) trained on P90 and tested 1 d later exhibit significantly less freezing (p = .0152) than Cre- control male mice (n = 5). No significant differences were determined during development: B Trained on P23 and tested 1 d later (p = .7501;  Cre-: n = 8; Cre+: n = 16) C Trained on P23 and tested 7 d later (p = .4680; Cre-: n = 6; Cre+: n = 12) D Trained on P35 and tested 30 d later (p = .5077; Cre-: n = 7; Cre+: n = 8) or E Trained on P50 and tested 30 d later (p = .6277; Cre-: n = 9; Cre+: n = 10). F There is no baseline freezing on conditioning day in P90 female control (n = 12) or cKO (n = 8) mice.
